# Cancer as a tissue anomaly: classifying tumor transcriptomes based only on healthy data

**DOI:** 10.1101/426395

**Authors:** Thomas P. Quinn, Thin Nguyen, Samuel C. Lee, Svetha Venkatesh

## Abstract

Since the turn of the century, researchers have sought to diagnose cancer based on gene expression signatures measured from the blood or biopsy as biomarkers. This task, known as classification, is typically solved using a suite of algorithms that learn a mathematical rule capable of discriminating one group (e.g., cases) from another (e.g., controls). However, discriminatory methods can only identify cancerous samples that resemble those that the algorithm already saw during training. As such, we argue that discriminatory methods are fundamentally ill-suited for the classification of cancer: because the possibility space of cancer is definitively large, the existence of a one-of-a-kind gene expression signature becomes very likely. Instead, we propose using an established surveillance method that detects anomalous samples based on their deviation from a learned normal steady-state structure. By transferring this method to transcriptomic data, we can create an anomaly detector for tissue transcriptomes, a “tissue detector”, that is capable of identifying cancer without ever seeing a single cancer example. Using models trained on normal GTEx samples, we show that our “tissue detector” can accurately classify TCGA samples as normal or cancerous and that its performance is further improved by including more normal samples in the training set. We conclude this report by emphasizing the conceptual advantages of anomaly detection and by highlighting future directions for this field of study.

## 1 Introduction

Cancer is a collection of complex heterogeneous diseases with known genetic and environmental risk factors. Physicians diagnose cancer by carefully weighing evidence collected from patient history, physical examination, laboratory testing, clinical imaging, and biopsy. Computers can aid diagnosis and improve outcomes by mitigating diagnostic errors. Indeed, this is already actively researched, where studies have shown that computers can reduce the reading errors of mammography [13] and commuted tomographic (CT) [3] images. Meanwhile, researchers have also sought to use computers to diagnose cancer based on gene expression signatures measured by high-throughput assays like microarray or next-generation sequencing [1, 4]. Gene expression signatures are ideal biomarkers because mRNA expression is dynamically altered in response to changes in the cellular environment. However, developing molecular diagnostics requires large data sets which have only recently become available due to reduced assay costs. These data could usher in a new era in clinical diagnostics.

Within the last decade, scientists have produced large transcriptomic data sets containing thousands of clinical samples. Of these, the TCGA stands out as the most comprehensive, having sequenced more than 10,000 unique tissue samples from 33 cancers and healthy tissue controls [19]. Meanwhile, an equally large study, GTEx, has sequenced non-cancerous samples comprising 54 unique human tissue types [10]. Already, a number of studies have used the TCGA data to build diagnostic classifiers that can determine whether a tissue sample is cancerous or not based only on its gene expression signature [7]. This task, known as classification, is typically solved using a suite of algorithms that learn a mathematical rule capable of discriminating one group (e.g., cases) from another (e.g., controls). This rule is learned from a large portion of the data called the “training set”, then evaluated on a withheld portion of the data called the “test set”. Discriminatory classifiers like artificial neural networks (ANNs), support vector machines (SVMs), and random forests (RFs) have become popular in the biological sciences [6]. All of these work well for high-dimensional data provided that the training set contains enough correctly labeled cases and controls.

In practice, clinicians need to answer questions like, "Is this tissue cancerous or not?" or "Is this cancer malignant or not?". ANNs, SVMs, and RFs can all answer these questions by learning a discriminatory rule from labeled data. However, discriminative methods have two major limitations, both of which apply to cancer classification. The first limitation is theoretical: discriminative methods suffer from the problem of having to see all possible abnormalities in order to make an accurate and generalizable prediction [15]. This is relevant to cancer because there exists countless ways in which a normal cell could become cancerous. As such, the label “cancer” does not encompass a known homogeneous group, but rather a heterogeneous collection of unknown types. It is simply not possible to anticipate the nature or extent of these “unknown unknowns” [14]. The second limitation is practical: even for a theoretically homogeneous cancer class, the tumor may occur too rarely for there to exist enough samples to inform a meaningful discrimination rule. Discriminatory methods require sufficient sample sizes to learn a rule that tolerates the large variance observed in replicates of transcriptomic data [11]. For these reasons, discriminatory methods are doomed to fail.

On the other hand, we expect that the possibility space for steady-state normal tissue is appreciably smaller than that of the aberrant tumor. By modeling this normal latent structure directly, we could learn a new rule that detects cancerous samples as a departure from normal. This follows the biological intuition that tumors themselves are anomalies of normal cellular physiology. The field of machine learning already has well-established methods that can detect anomalies in high-dimensional data, especially images, for the purpose of surveillance [2]. By transferring these methods to transcriptomic data, we can create an anomaly detector for tissue transcriptomes, a “tissue detector”, that is capable of identifying cancer without ever seeing a single cancer example. In this report, we show that “tissue detectors” are sensible and accurate for the classification of cancer based on gene expression signatures. We do this by training an anomaly detection model on normal GTEx samples, then using it to accurately differentiate normal from cancerous TCGA samples. In presenting these results, we highlight future research directions for the detection of anomalous gene expression signatures.

## 2 Methods

We acquired the combined GTEx and TCGA data from Wang et al. 2018, who harmonized the disparate data sets using quantile normalization and svaseq-based batch effect removal [18]. The Wang et al. data represents six tissues as characterized in Table 1: breast, liver, lung, prostate, stomach, and thyroid. We accessed the data as fragments per kilobase of transcript per million (FPKM), separated the data by tissue, and z-score standardized each gene. We then performed a residual analysis on each tissue with GTEx training sets and TCGA test sets. Residual analysis is based on the principle that most data have an underlying structure that can be largely reconstructed using a subset of the principal components, whereby the difference between the reduced representation and the original observations are termed the *residues*. Residual analysis uses the squared value of the residue as a proven way to measure the degree to which an observation is an outlier. For normally distributed data, the squared value of the residues follows a non-central *χ*^2^ distribution. By comparing the norm of the residue for an unlabeled sample to a procedurally generated threshold (corresponding to a stipulated false alarm rate), we have a predictive rule that decides whether to reject the null hypothesis and call that sample an anomaly [5].

We refer to a predictive model and its threshold as a “tissue detector”, of which we trained six (one for each tissue). After training each model on the GTEx data, we evaluated its performance on the respective TCGA data. For each sample in the test set, we calculated an anomaly score based on the distance between that sample and the model reference. We do this by projecting the sample to the principal component space and measuring its residue, where higher residue scores indicate that the sample is more anomalous. If the anomaly score is larger than the anomaly detection threshold, the sample is called abnormal (i.e., an outlier). Otherwise, the sample is called normal (i.e., an inlier). This allows us to differentiate between normal and cancerous TCGA samples without ever seeing a single cancer example. We repeated this procedure for increasingly smaller subsets of the training data, with specificity averaged across ten bootstraps each. We present these results in Figure 2.

## 3 Results

In this study, we trained a “tissue detector” on each six tissues represented in the combined GTEx and TCGA data set, using only the GTEx samples for training. We then evaluated its performance on the withheld TCGA data by calculating an anomaly score for each TCGA sample and comparing it against the anomaly threshold: if the score is greater than the threshold, the sample is considered an anomaly (i.e., cancerous). Figure 1 shows the (log-)ratio of per-sample anomaly scores relative to the tissue-specific anomaly threshold (y-axis) for each tissue (x-axis), faceted based on whether the sample is cancerous. Especially for breast, liver, lung, and thyroid data, our “tissue detector” not only recognizes most TCGA cancer samples as anomalies, but also recognizes most TCGA healthy samples as normal. On the other hand, anomaly detection performs poorly for prostate and stomach tissue. Table 1 shows the precision, recall, and specificity for each “tissue detector”.

**Figure 1:**
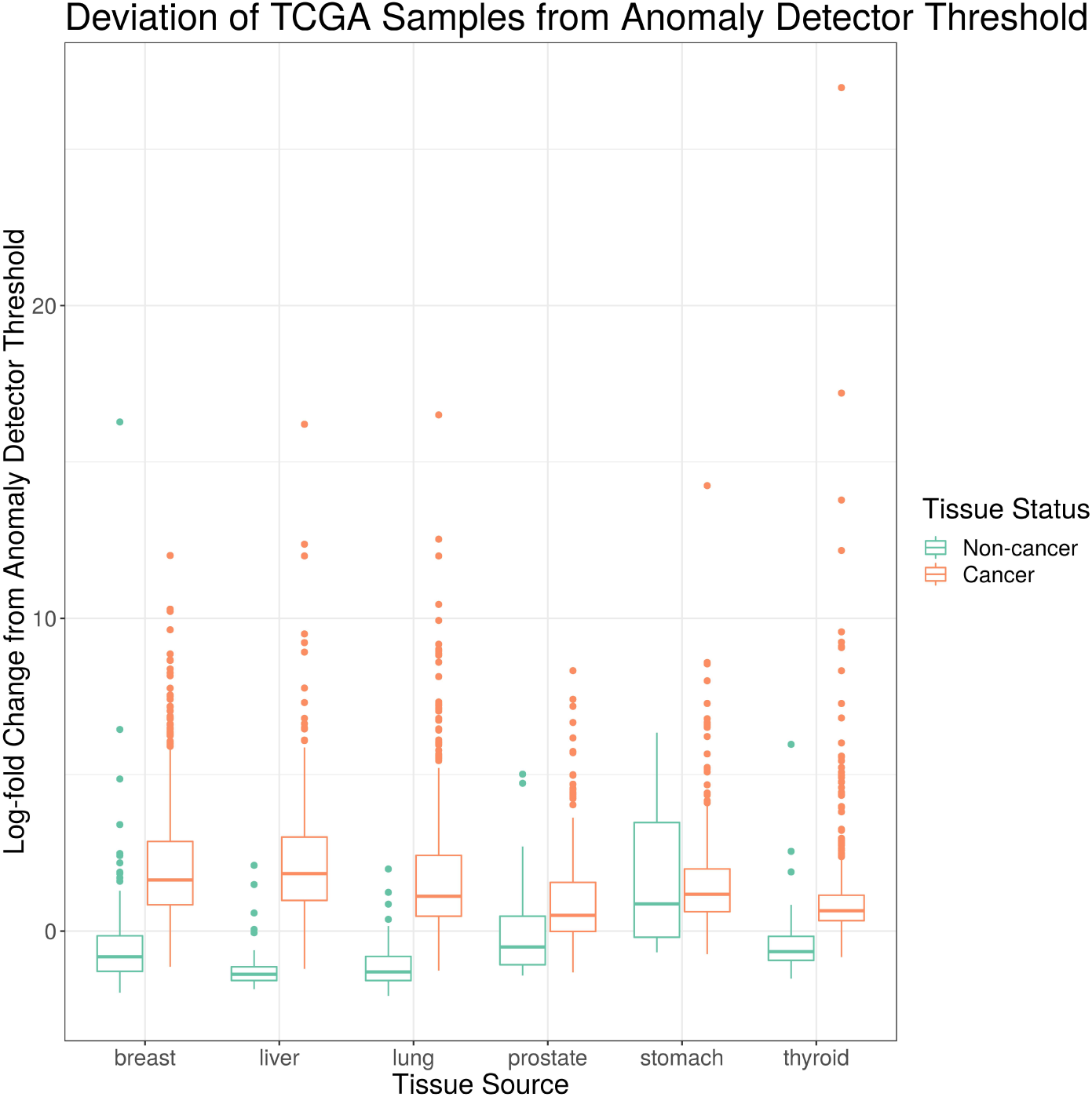
This figure shows the (log-)ratio of per-sample anomaly scores relative to the tissue-specific anomaly threshold (y-axis) for each tissue (x-axis), faceted based on whether the sample is cancerous. The “tissue detector” calls any sample above the x-intercept threshold as an anomaly (i.e., cancerous). The threshold is determined procedurally during model training. This figure shows performance for TCGA test set only; no TCGA samples were included in the training set.

**Table 1:** This table shows the number of samples in each GTEx training set and TCGA test set, alongside the test set performance of that anomaly detector. Precision and recall remain high for all classifiers, but specificity suffers for select tissues. This suggests that our “tissue detector”, when it fails, has a bias toward viewing all TCGA samples as abnormal. The acronyms N and C refer to number of normal and cancerous samples, respectively.

|  | GTEX (N) | TCGA (N) | TCGA (C) | Precision | Recall | Specificity | Accuracy | AUC |
| --- | --- | --- | --- | --- | --- | --- | --- | --- |
| breast | 89 | 110 | 982 | 0.975 | 0.965 | 0.782 | 0.947 | 0.903 |
| liver | 115 | 48 | 295 | 0.986 | 0.939 | 0.917 | 0.936 | 0.973 |
| lung | 313 | 59 | 503 | 0.987 | 0.907 | 0.898 | 0.906 | 0.960 |
| prostate | 106 | 48 | 426 | 0.949 | 0.742 | 0.646 | 0.732 | 0.734 |
| stomach | 192 | 33 | 380 | 0.943 | 0.966 | 0.333 | 0.915 | 0.547 |
| thyroid | 318 | 53 | 441 | 0.974 | 0.925 | 0.792 | 0.911 | 0.893 |

For all tissues, our anomaly detectors tended to have better sensitivity (i.e., recall) than specificity. Intuitively, we expect that increasing the number of normal samples shown to the “tissue detector” during model training would improve its specificity, especially for the poorly performing prostate and stomach detectors. To test this hypothesis, we measured the specificity of each “tissue detector” as trained on increasingly smaller subsets of the GTEx data. Figure 2 shows the specificity for each “tissue detector’ (y-axis) according to the number of samples in the training set (x-axis). A pattern emerges: the inclusion of additional GTEx samples can improve the classification of TCGA samples, up until a point of diminishing returns.

**Figure 2:**
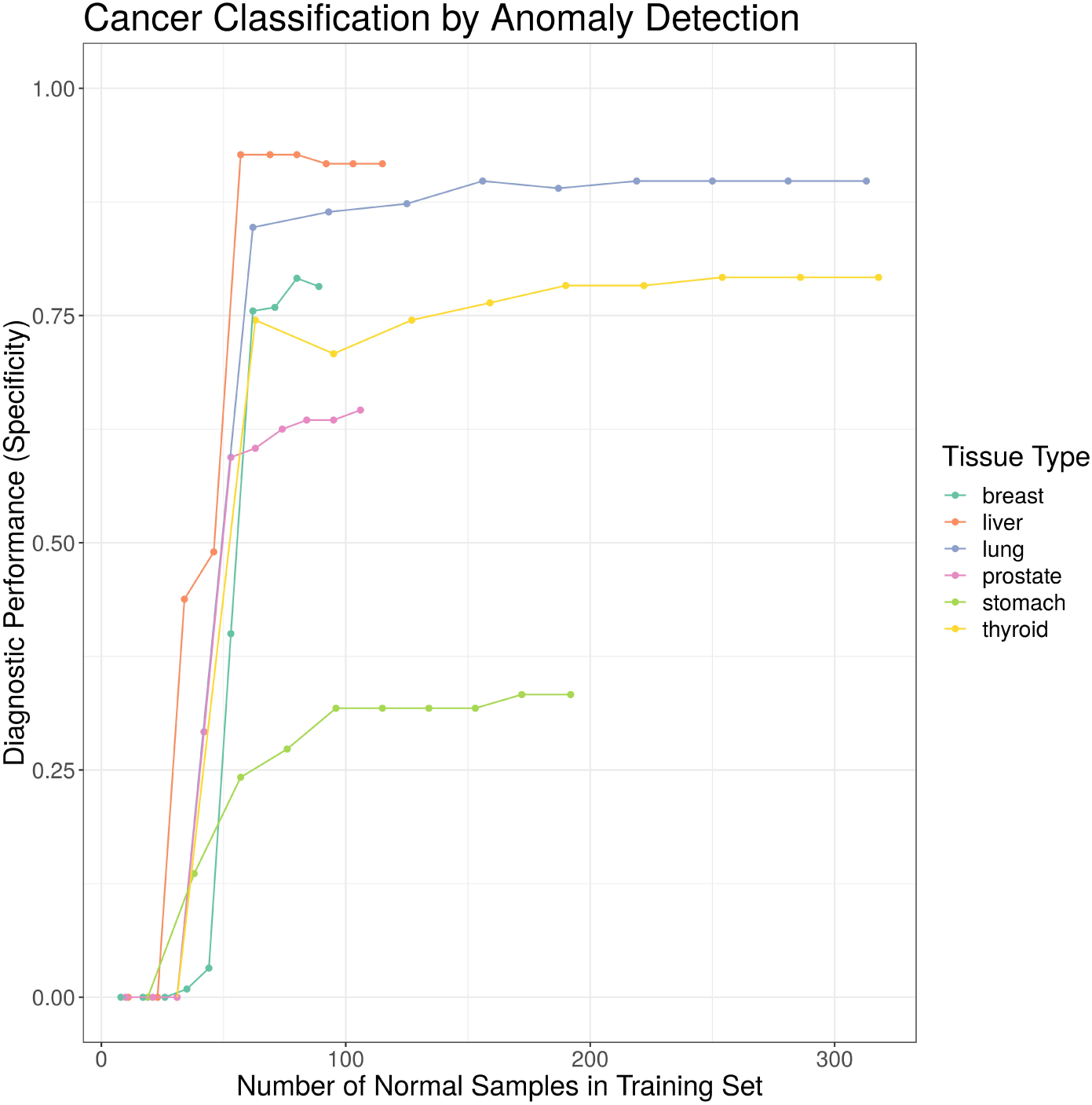
This figure shows the specificity for each “tissue detector” (y-axis) according to the number of samples in the training set (x-axis). Performance is averaged across 10 bootstraps of the GTEx training set. This figure shows performance for TCGA test set only; no TCGA samples were included in the training set.

## 4 Discussion

Technological advances have made it possible to measure the global gene expression signature of any biological sample at little cost. Already, there is a growing body of evidence that gene expression signatures can be used as biomarkers to diagnose cancer [7]. In this report, we present a novel application of anomaly detection to classify cancer based on gene expression signatures. By learning the latent structure of normal gene expression from a training set of normal samples, we created a “tissue detector” that can identify cancer without having seen a single cancer example. Our method contrasts with discriminatory methods, widely used in the biological sciences, which can only identify cancerous samples that resemble those that the algorithm already saw during training. In principle, discriminatory methods do not make sense for a disease like cancer where a one-of-a-kind gene expression signature is theoretically possible. Practically speaking, anomaly detection further benefits from normal samples being more readily available and easier to collect than abnormal samples: for any cancer, many more people do not have the cancer than do. Since the inclusion of additional normal samples can improve the specificity of anomaly detection, as demonstrated in Figure 2, the curation of large normal data sets could open up the possibility of building diagnostic tests for extremely rare cancers.

Although we applied anomaly detection here to differentiate normal from cancerous tissue, anomaly detection could suit a number of other health surveillance applications. By changing the class of samples used in the training set, the meaning of “anomaly” changes. For example, if we include only benign tumors in the training set, then an anomaly detector might identify whether a biopsied tumor is potentially malignant (i.e., not benign). Likewise, using a training set of blood biomarkers for patients with surgically resected tumors might yield an anomaly detector that can identify whether a primary tumor has recurred. Other novel applications might include training a “tissue detector” on healthy lymphatic tissue to screen for lymphatic metastasis or on chemotherapy-sensitive tumor biopsies to screen for emerging drug resistance. Whatever the application, anomaly detection is unique in that it only requires that there exist data for the null state that is under surveillance: it is not necessary that researchers have characterized the full spectrum of the undesired outcome.

One challenge faced during the detection of anomalous gene expression signatures is the limited amount of data available for training and testing. Even as data sets get larger, anomaly detection will still benefit from the combination of multiple data sets, known as horizontal data integration [17]. However, horizontal data integration is complicated because every data set has intra-batch and inter-batch effects caused by systematic or random differences in sample collection. These differences could arise from a variety of biological factors (e.g., biopsy site, age, sex) or technical factors (e.g., RNA extraction protocol, sequencing assay), including latent factors unknown to the investigator [9]. Although software like ComBat and sva can remove intra-batch biases, inter-batch biases may still remain. Indeed, inter-batch biases could explain why our “tissue detectors”, when they fail, tend to view all TCGA samples as abnormal (though the “normal” TCGA samples do all come from sites adjacent to cancerous tissue). Although Wang et al. tried to harmonize the TCGA and GTEx data [18], the removal of inter-batch biases is non-trivial and further challenged by the prevailing need to preserve test set independence. Moreover, owing to how next-generation sequencing data measure the relative abundance of gene expression, these data also contain inter-sample biases that sit on top of the intra-batch and inter-batch biases [16, 12]. It remains an open question of how best to integrate multiple data sets. Non-parametric or compositional PCA-like methods could provide a suitable alternative to anomaly detection that is more robust to inter-batch and inter-sample biases.

Another challenge faced during the detection of anomalous gene expression signatures is the lack of transparency in the decision-making process. Although the concept of anomaly detection is intuitive, its implementation decomposes high-dimensional data into orthogonal eigenvectors that do not necessarily have any meaning to biologists. When examining these eigenvectors directly, it may be unclear how an anomaly detection model reached its decision. This makes it difficult to formulate new hypotheses to improve the model performance or elucidate the biological system. Future work should aim to improve the interpretability of anomaly detection methods. One approach might involve building a tool that visualizes which eigenvector components contributed maximally to each decision. If some constituent genes are consistently involved in misclassification, this could generate testable hypotheses. Similarly, one could try to characterize the biological importance of the maximally relevant eigenvectors through gene set enrichment analysis (GSEA), as done by Weighted Gene Correlation Network Analysis [8]. This would allow investigators to frame inlier and outlier distributions not only in terms of the constituent genes involved, but also in terms the biological pathways affected. This too could generate testable hypotheses. With these improvements, anomaly detection would become an interpretable and actionable classification strategy for many health surveillance applications.

